# Influence of different modes of morphological character correlation on phylogenetic tree inference

**DOI:** 10.1101/308742

**Authors:** Thomas Guillerme, Martin D. Brazeau

## Abstract

Phylogenetic analysis algorithms require the assumption of character independence - a condition generally acknowledged to be violated by morphological data. Correlation between characters can originate from intra-organismal features, shared phylogenetic history or forced by particular character-state coding schemes. Although the two first sources can be investigated by biologists *a posteriori* and the third one can be avoided *a priori* with good practices, phylogenetic software do not distinguish between any of them.

In this study, we propose a new metric of raw character difference as a proxy for character correlation. Using thorough simulations, we test the effect of increasing or decreasing character differences on tree topology. Overall, we found an expected positive effect of reducing character correlations on recovering the correct topology. However, this effect is less important for matrices with a small number of taxa (25 in our simulations) where reducing character correlation is not more effective than randomly drawing characters. Furthermore, in bigger matrices (350 characters), there is a strong effect of the inference method with Bayesian trees being consistently less affected by character correlation than maximum parsimony trees.

These results suggest that ignoring the problem of character correlation or independence can often impact topology in phylogenetic analysis. However, encouragingly, they also suggest that, unless correlation is actively maximised or minimised, probabilistic methods can easily accommodate for a random correlation between characters.

## Introduction

The last two decades have witnessed a “resurgence” of interest in the use of morphological character data in phylogenetic studies. This owes in large part to the use of fossils to undertake at least partial reconstructions of phylogenetic trees, especially where ancestral states reconstructions or absolute calibrations of divergence times are necessary. Morphological character data are often considered inferior to molecular sequence data, but are often the only source of phylogenetic data for extinct species. While there is a general appreciation of the limits of morphological data, they are frequently dismissed without any empirical investigations into their statistical properties. As morphological data are likely to continue to play an extensive role in phylogenetic analysis, it is essential to understand the circumstances under which morphological data might be expected to “misbehave”. This opens up possibilities for predicting problematic datasets and possibly proposing new confidence measures in phylogenetic datasets.

The non-independence of large numbers of morphological characters is often cited in anticipation of problems with morphological data. The assumption of character independence is central to phylogenetic inference methods such as maximum likelihood and maximum parsimony (e.g. Joysey and Friday, 1982; Felsenstein, 1985; Lewis, 2001; Felsenstein, 2004). Especially for discrete morphological data, this assumption of independence is probably violated frequently due to the very nature of phylogenetic data: correlations are expected to occur (to some degree) when characters are depending on each other. Before discussing character correlation further, it is important to understand that it may manifest itself in at least three distinct ways:

- **Intra-organismal dependence:** this is the result of an intrinsic biological link between two characters through development, pleiotropy, and/or biological function. For example the lower and upper molar characters in mammals generally occlude one another. Therefore, one character describing a feature of a lower molar will be expected to be complemented by the surface of the occluding upper molar. Characters of the occlusal surface of two opposing molars will be expected to directly covary. Pleiotropy also results in covariation between different aspects of phenotype. From a phylogenetic perspective, it can be especially pernicious because the relationship between the traits in question may have no obvious link from a morphological or functional comparison alone. Intra-organismal links can be the targets of comparative developmental biology (Goswami and David Polly, 2010; Kelly and Sears, 2010; Stoessel et al., 2013; Goswami et al., 2014) or functional investigations.
- **Evolutionary dependence:** this is the result of sets of characters co-evolving due to selection, likely related to functional links between two traits that help serve an overall lifestyle trait. Unlike the case of intra-organismal dependence, there need not be an intrinsic constraint that causes these traits to covary. For example, in vertebrates, axial elongation can be correlated to limb reduction with snake-like bodies evolving multiple times in numerous tetrapod lineages. This is thought to correspond to adaptations for fossoriality or aquatic lifestyles. Such covariances are generally studied in the context of a given phylogeny, often one derived from molecular data with the morphological traits of interest mapped on it. Many methods have been developed to study these correlations, especially since they can provide us with a lot of information on how specific groups acquired specific characteristics (Russell Lande, 1983; Maddison, 1990; Pagel, 1994; Mark Pagel, 2006; Grabowski and Porto, 2016). However, these methods do not give us a means to objectively control correlations that might adversely affect phylogenetic inference.
- **Coding dependence:** this is the results of researcher methodology for defining or/and coding discrete morphological characters (Brazeau, 2011; Simôes et al., 2017). Coding dependence manifests itself in several ways, particularly in coding redundant information. For instance, coding for the same absence in different characters creates state transformations associated with the loss or gain of a particular character. This occurs when a number of multistate characters include two variable feature states (e.g. large, small; red, blue etc.) in conjunction with absence. It is worth noting, however, that these correlations could also be due to the nature of the available data, especially in palaeontology. For example, when only one fragmentary molar is available to describe a specimen, researchers have to “extract” as much phylogenetic information from the available data as possible, potentially inducing correlations. This coding dependency is linked to hierarchical dependency between characters (Wilkinson, 1995; Brazeau et al., 2017). Finally this can also be due to a bias in the amount of characters available. For example, in skulls, because of their complexity, there is a high likelihood of inducing correlation (by effectively reducing structural complexity to discrete characters).

Of course, the three sources of dependence have an interaction: characters describing the left and right lower/upper molars will have induced dependence due to the modularity of the molars, their shared history and the duplicated coding. Logical dependence, however, is easily distinguished prior to phylogenetic inference, while the two other ones (intra-organismal and evolutionary) are much harder. However, the development of algorithms and software has not yet caught up with the need to deal with these interdependencies (De Laet, 2015; Brazeau et al., 2017). Intra-organismal dependence requires more detailed, often extremely time-consuming studies (and possibly beyond the limits of available technology). Even after all of the effort is expended, the results might then only be known for a single (model) species. Evolutionary dependence itself requires the resolution of a phylogenetic tree, and is best determined by independent character sets. This is frequently accomplished by mapping morphological traits on molecular phylogenetic trees.

These sources of dependence between characters are well studied in biology. Biological and evolutionary dependences are inherent parts to Evo-Devo and macroevolutionary studies and best practices to avoid coding-induced dependences are commonly known and applied. However, eventually, all these characters, whether they are independent or not are analysed through phylogenetic inferences software that are blind to these distinctions. If fact, what the software are confronted with is a two dimensional matrix problem that renders the morphological subtleties described opaque. This introduces a new, less studied, source of character dependence: **Correlation between characters detected by the software**: this is the result how software actually interprets the differences between characters. The vast majority of phylogenetic software ignores both the character’s definition and the different states signification (simply treating them as different or similar tokens). Therefore a great number of characters and - traditionally - a few number of tokens can easily lead to dependence between characters. For example, if we consider the following matrix containing four cetartiodactyls - say a pig (e.g. *Sus*), a deer (*Cervus*), a hippo (*Hippopotamus*) and a whale (*Balaenoptera*) - and four binary characters - say (**C_1_**: presence (1) or absence (0) of an astragalus; **C_2_**: presence (0) or absence (1) of baleen; **C_3_**: presence (0) or absence (1) of a left astragalus with a double pulley; **C_4_**: presence (0) or absence (1) of a right astragalus with a double pulley:

In the example in Table 1, the characters **C_1_** and **C_2_** are the most likely to be truly independent; characters **C_3_** and **C_4_** suffer from a a coding induced dependency; characters **C_1_** and **C_3_**/**C_4_** have an evolutionary induce dependency and again, characters **C_2_** and **C_3_**/**C_4_** are likely to be independent. Yet a phylogenetic software will treat all these four characters in exactly the same way: only the sheer difference between the character states tokens will be used in order to infer the tree. Some characters will therefore be expected to covary in non-phylogenetic way, and that this phenomenon can reasonably be expected to mislead phylogenetic analysis. Yet the question has never been explored through a thorough simulation framework (although it has been tackled empirically for morphological data Dávalos et al. 2014 or molecular data Zou and Zhang 2016).

**Table 1:** Example of a matrix with software induced character correlation. **C_1_**: presence (1) or absence (0) of an astragal; **C_2_**: presence (0) or absence (1) of baleens; **C_3_**: presence (0) or absence (1) of a left astragalus with a double pulley; **C_4_**: presence (0) or absence (1) of a right astragal with a double pulley.

|  | C1 | C2 | C3 | C4 |
| --- | --- | --- | --- | --- |
| *Sus* | 1 | 1 | 1 | 1 |
| *Cervus* | 1 | 1 | 1 | 1 |
| *Hippopotamus* | 1 | 1 | 1 | 1 |
| *Balaenoptera* | 0 | 0 | 0 | 0 |

How does these correlation really affect topology? We expect matrices with a high level of correlation to recover precise but inaccurate topologies but will matrices with low level of correlation (i.e. with high levels of homoplasy) actual cancel out the effects of correlation? Here we formally assess the effect of discrete character’s correlation using simulated data. We propose a new distance metric to measure the difference between characters (as a proxy for these three sources of correlation as interpreted by the software) and a protocol to modify discrete morphological matrices to increase/decrease the overall differences or similarities between characters. We found that overall, there is a detectable effect of character correlation on topology where an increase in character dependence results in a decrease in the ability to recover the correct topology. These results, however, vary greatly in magnitude depending on the size of matrix and the inference method used.

## Methods

To assess the effects of character correlation on the accuracy of phylogenetic inference we generated a series of matrices exhibiting different levels of correlation between some characters (Fig.1 - note that each step is described in more details below):

1. **Simulating matrices**: we simulated discrete morphological matrices with 25, 75 and 150 taxa for 100, 350 and 1000 characters, hereafter called the “normal” matrices. This step resulted in 9 matrices.
2. **Modifying matrices**: we changed the “normal” matrices by modifying the characters in order to maximise or minimise characters differences (hereafter called respectively “maximised” and “minimised” matrices) by removing respectively the least different or most different characters and replacing them randomly by the remaining characters. Our protocol for measuring character difference is detailed below. We also randomly duplicated characters from the “normal” matrices without biasing towards maximising or minimising character differences to create randomised matrices (hereafter called the “randomised” matrices - equivalent to a null expectancy). This step resulted in 36 matrices.
3. **Inferring topologies**: we inferred the topologies from the “normal”, “maximised”, “minimised” and “randomised” matrices using both maximum parsimony and Bayesian inference. Hereafter, the resultant topologies are called the “normal”, “maximised”, “minimised” and “randomised” trees). This step resulted in 72 topologies.
4. **Comparing topologies**: finally, we compared the “normal” to the “maximised”, “minimised” and “randomised” trees to measure the effect of character correlation on topology.

Each step was replicated 35 times and are described below in more detail, along with our proposed definition for measuring the difference between characters.

**Figure 1:**
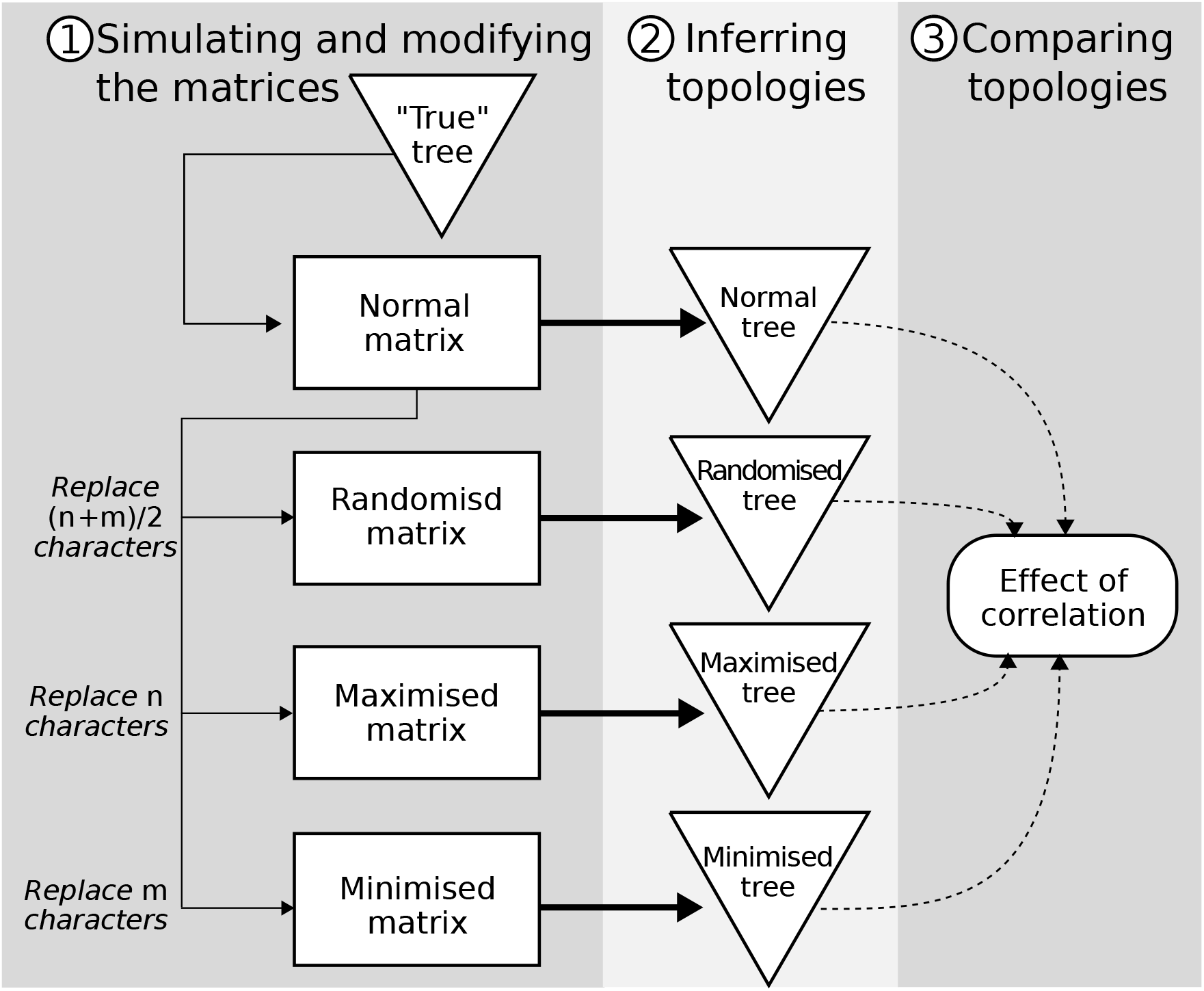
Outline of the simulation protocol: the first step includes both the simulation and the modification of the matrices (thin solid lines); the second step includes tree inference using MP and BPP methods (thick solid lines); the third step includes comparing the resulting tree topologies (dashed lines). *n* and *m* corresponds to the number of characters with a character difference < 0.25 and > 0.75 respectively.

### Measuring differences between characters

To measure the effect of character correlation as interpreted by the phylogenetic software, we define characters as being entirely correlated if they give the same phylogenetic information. In order to measure this, we propose a new distance metric to measure the difference between two characters:

#### Character Difference (CD)

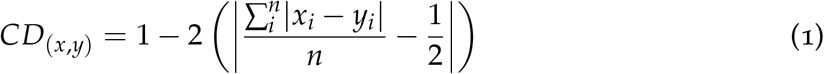

Where *n* is the number of taxa with comparable characters *x*, *y* and *x*_*i*_, *y*_*i*_ are each character’s state for the *i*^*th*^ taxon. *CD* is a continuous distance metric bounded between 0 and 1 (see the mathematical demonstration in the supplementary material 1). Since we are considering differences as being only Fitch-like (non-additive) and unweighted, we calculated the difference between character states in a qualitative way. Two same character states tokens have a difference of zero and two different ones have a difference of one (e.g. 0 − 0 = 0 or 1 − 8 = 1). Additionally, we only consider differences for taxa with shared information (i.e. a Gower distance; Gower, 1971).

We standardised each character by arbitrarily modifying their character state tokens (or symbols) by order of appearance. In other words, we replaced all the occurrences of the first token to be 1, the second to be 2, etc. This procedure was used to treat all the characters are unordered with no assumption on the meaning of the character state (e.g. in a binary character 0 is not necessary ancestral to 1). It also greatly improved the speed of our algorithm implementation to compare the characters. This way, a character `A = {2,2,3,0,0,3}` for six taxa would be standardised as `A’ = {1,1,2,3,3,2}` (following the *xyz* notation in Felsenstein, 2004, p.13). Note that in terms of phylogenetic signal, both `A` and `A’` are exactly identical (forming three distinct splits in the tree inference process).

When the character difference is null (0) it means that characters convey the same phylogenetic signal (i.e. characters are entirely correlated). When the character difference is maximal (1) it means it conveys the greatest difference in phylogenetic signal (i.e. characters are uncorrelated). It is important to stress that a character difference of 0 (i.e. the same phylogenetic signal) does not mean the opposite of 1 (i.e. *not* the opposite phylogenetic signal but the most different number of implied splits). For example with three characters `A = {0,1,1,1}`, `B = {1,0,0,0}` and `C = {0,1,2,3}`, *CD*_(*A*,*B*)_ = 0 and *CD*_(*A*,*C*)_ = 1. Because the character is continuous and bounded between (0, 1), it can be interpreted as the probability of two characters leading to a different set of splits (i.e. a different phylogenetic signal).

### Simulating discrete morphological matrices

To simulate the matrices we applied a protocol very similar to Guillerme and Cooper (2016b). First, we generate random birth-death trees with the birth (*λ*) and death (*μ*) parameters sampled from a uniform (0, 1) distribution maintaining *λ* > *μ* using the `diversitree R` package (v0.9-8; FitzJohn, 2012) and saving the tree after reaching either 25, 75 or 150 taxa. For each tree, we arbitrarily set the outgroup to be the first taxon (alphabetically) thus effectively rooting the trees on this taxon. These trees are hereafter called the “true” trees (see distinction below). We then simulated discrete morphological characters on the topology of these trees using the either of the two following models:

- The morphological HKY-binary model (O’Reilly et al., 2016) which is an HKY model (Hasegawa et al., 1985) with a random states frequency (sampled from a Dirichlet distribution *Dir*(1, 1, 1, 1)) and using a transition/transvertion rate of 2 (Douady et al., 2003) but where the purines (A,G) were changed into state 0 and the pyrimidines (C,T) in state 1. This model has the advantage of not favouring Bayesian inference (since it doesn’t use an M*k* model; O’Reilly et al., 2016,; see discussion) but the downside of it is it can only generate binary state characters (or 4 states; Puttick et al., 2017).
- To generate more than binary states characters, we used the M*k* model (Lewis, 2001). We draw the number of character states with a probability of 0.85 for binary characters and 0.15 for three state characters (Guillerme and Cooper, 2016b; Zou and Zhang, 2016). This model assumes a equal transition rate between character states which might seem overly simplistic, excluding other observed transition patterns (e.g. Dollo characters; Dollo, 1893; Wright et al., 2015). Recently however, Wright et al. (2016) have shown that an equal rate transition is still the most present in empirical data.

For each character, both models (morphological HKY-binary or M*k*) where chosen randomly and run with an overall evolutionary rate drawn from a gamma distribution (*β* = 100 and *α* = 5). These low evolutionary rate values allowed reduction in the number of homoplasic character changes, thus reinforcing the phylogenetic information in the matrices. We re-simulated every invariant characters to obtain a matrix with no invariant characters in order to better approximate real morphological data matrices. To ensure that our simulations were reflecting realistic observed parameters, we only selected matrices with Consistency Indices (CI) superior to 0.26 (O’Reilly et al., 2016).

For each tree with 25, 75 or 150 taxa we generated matrices with 100, 350 and 1000 characters following O’Reilly et al. (2016). The matrices were generated using the `dispRity R` package (Guillerme, 2016). To estimate the variance of our simulations and assess the effect of our random parameters, we repeated this step 35 times resulting in 315 “normal” morphological matrices.

### Modifying the matrices

We calculated the pairwise character differences for each generated matrix using the `dispRity R` package (Guillerme, 2016). We then modified the matrices to either maximise or minimise the pairwise character differences for each matrix using three different algorithms. For maximising the pairwise differences between characters, we selected the characters that were the most similar to all the others (i.e. with an average character difference < 0.25) and replaced them randomly by any of the remaining characters. This operation increased the overall pairwise character difference in the matrix thus making the characters more dissimilar. Conversely, for minimising the pairwise character differences, we selected the most dissimilar characters (i.e. with an average character difference < 0.75) and randomly replaced them with the remaining ones. Finally, because this operation effectively changes the weight of characters (i.e. giving the characters < 0.25 or > 0.75 a weight of 0 and giving the randomly selected remaining characters a weight of +1), we randomly replaced the average number of characters replaced in the character maximisation and minimisation by any other characters as a randomised expectation modification (i.e. randomly weighting characters). Each of the three matrices are effectively a bootstrap pseudo-replication of the “normal” matrix with the “randomised” one being a random one and the “maximised” and “minimised” being conditional bootstraps. This step resulted in a total of 1260 matrices (hereafter called the “normal”, “maximised”, “minimised” and “randomised” matrices - see Fig. 2 for an illustration). The algorithms for the three modifications are available on GitHub (https://github.com/TGuillerme/CharactersCorrelation)

**Figure 2:**
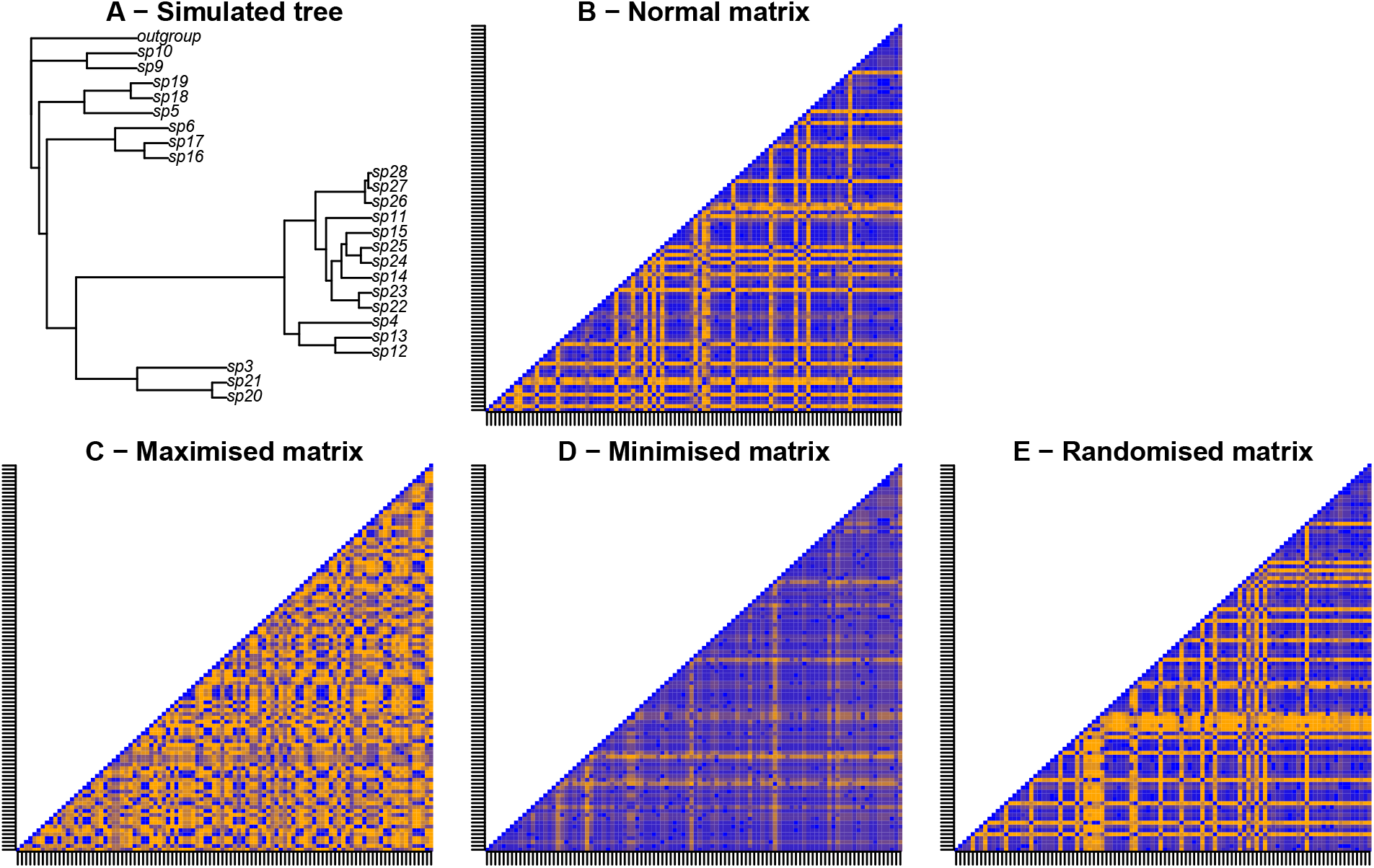
Example illustration of the protocol for modifying matrices. The matrices represent the pairwise character differences for 100 characters. Blue colours correspond to low character differences and orange colours correspond to high character differences. **A** - a random Birth-Death tree is simulated and used for generating the “normal” matrix (**B**), characters in this matrix are then removed or duplicated to favour maximised (**C**), minimised (**D**) or randomise character difference (**E**). The differences between the characters is low in **C** (minimised compared to **A**) implying a high correlation between the characters. Conversely, the character differences is high in **D** (maximised compared to **A**) implying a low correlation between the characters.

### Inferring topologies

We inferred the topologies with both BPP and MP using MrBayes (v3.2.6; Ronquist et al., 2012) and PAUP* (v4.0a151; Swofford, 2001) respectively. For both methods, we used the arbitrarily chosen outgroup in the simulations to root our trees. The maximum parsimony inference was run using a heuristic search with random sequence addition replicate 100 times with a limit of 5 × 10^6^ rearrangements per replicates (`hsearch addseq=random nreps=100 rearrlimit=5000000 limitperrep=yes`).

Bayesian inference was run using an M*k* model with ascertainment bias and four discrete gamma rate categories (M*kv* 4G - `lset nst=1 rates=gamma Ngammacat=4`) with an variable rate prior an exponential (0.5) shape (`prset ratepr=variable Shapepr=Exponential(0.5)`). We ran two runs of 6 chains each (2 hot, 4 cold) for a maximum of 1 × 10^9^ generations with a sampling every 200 generations. We automatically stopped the MCMC when the average standard deviation of split frequencies (ASDSF) between both runs fell below 0.01 (with a diagnosis every 1 × 10^4^ generations - `mcmc nruns=2 Nchains=6 ngen=1000000000 samplefreq=200 printfreq=2000 diagnfreq=10000 Stoprule=YES stopval=0.01 mcmcdiagn=YES`). Due to cluster hardware requirements an to save some time, when chains didn’t converged and the runs exceeded 5GB each, we aborted the MCMC and computed the consensus tree from the unconverged chains. In practice, these few MCMC got stuck at an ASDSF around (but not below) 0.01.

A strict majority rule tree was then calculated for both Bayesian an maximum parsimony trees. For the Bayesian consensus trees, the 25% first trees of the posterior tree distribution were excluded as a burnin. The 2880 tree inferences took around one CPU century on the Imperial College High Performance Computing Service (2-3GHz clock rate; ICHPC, 2011).

### Comparing topologies

We compared the topologies using the same approach as in Guillerme and Cooper (2016b): we measured both the Robinson-Foulds distance (Robinson and Foulds, 1981) and the triplets distance (Dobson, 1975) between the trees inferred from the “maximised”, “minimised” and “randomised” matrices and the tree inferred from the “normal” matrix. We explored the effect of character difference on recovering the “normal” topology by comparing the “maximised”, “minimised” and “randomised” trees to the “normal” tree (Figs 3 and 4 and supplementary materials 3 Figs 1 and 2). Note that we are not comparing the trees to the “true” tree used to simulate the matrices. First, in biology, this tree is always unknown. Second, our objective is to measure the direct effect of character correlation approximated by the difference in topology between the “normal”, “maximised” and “minimised” trees. When measuring the difference between these trees and the “true” tree, we would also confound the effect of simulating a birth-death tree and simulating a discrete morphological matrices from it.

**Figure 3:**
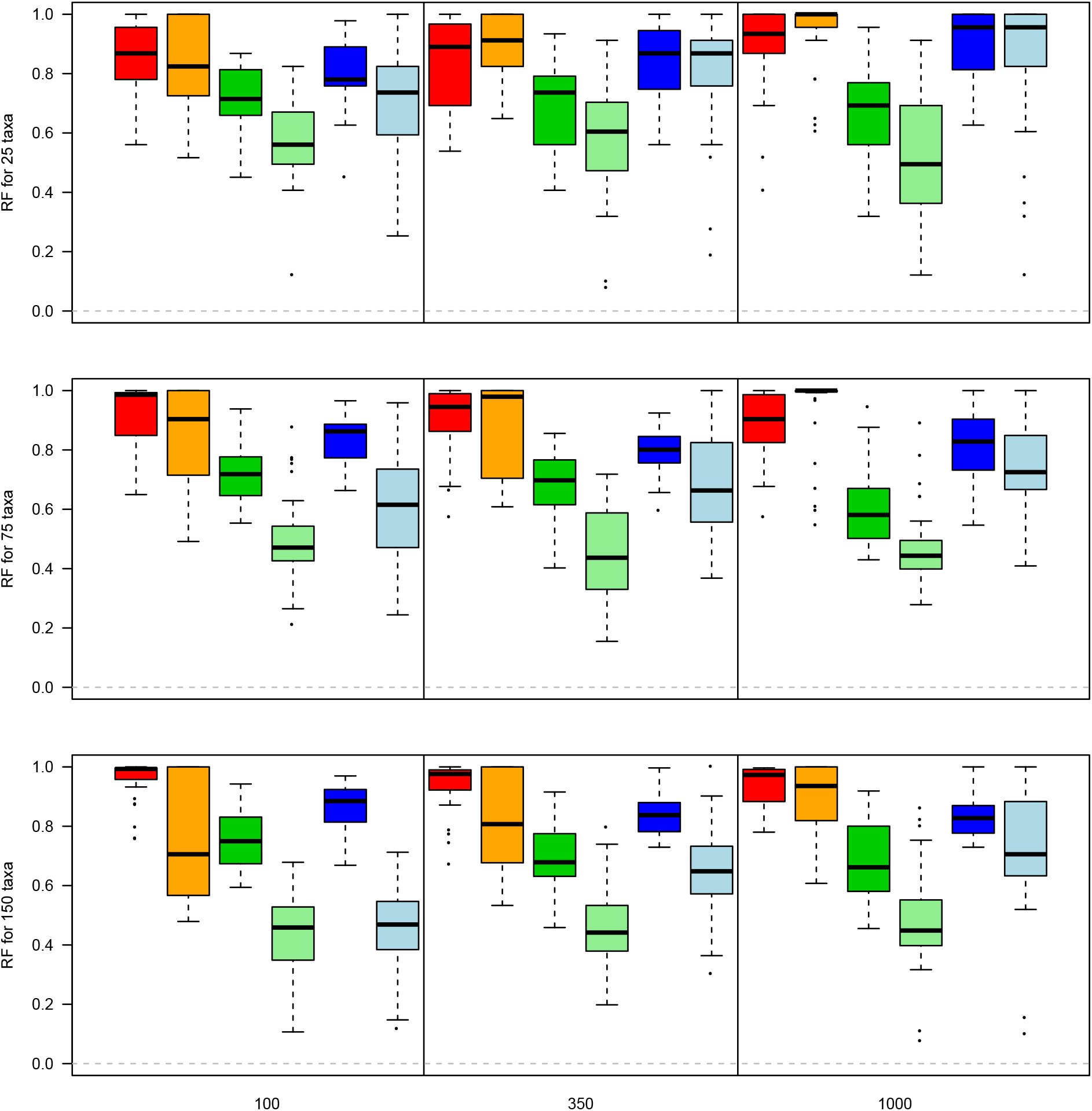
Effect of character difference on recovering the “normal” topology. The y axis represents the Normalised Tree Similarity using Robinson-Fould distance for matrices with 25, 75 and 150 taxa from top to bottom respectively. The x axis represents the different character difference scenarios and tree inference method with the “maximised” character difference in Bayesian (red) and under maximum parsimony (orange), the “minimised” character difference in Bayesian (dark green) and under maximum parsimony (light green) and the “randomised” character difference in Bayesian (dark blue) and under maximum parsimony (light blue) for matrices of 100, 350 and 1000 characters in the panels from left to right.

**Figure 4:**
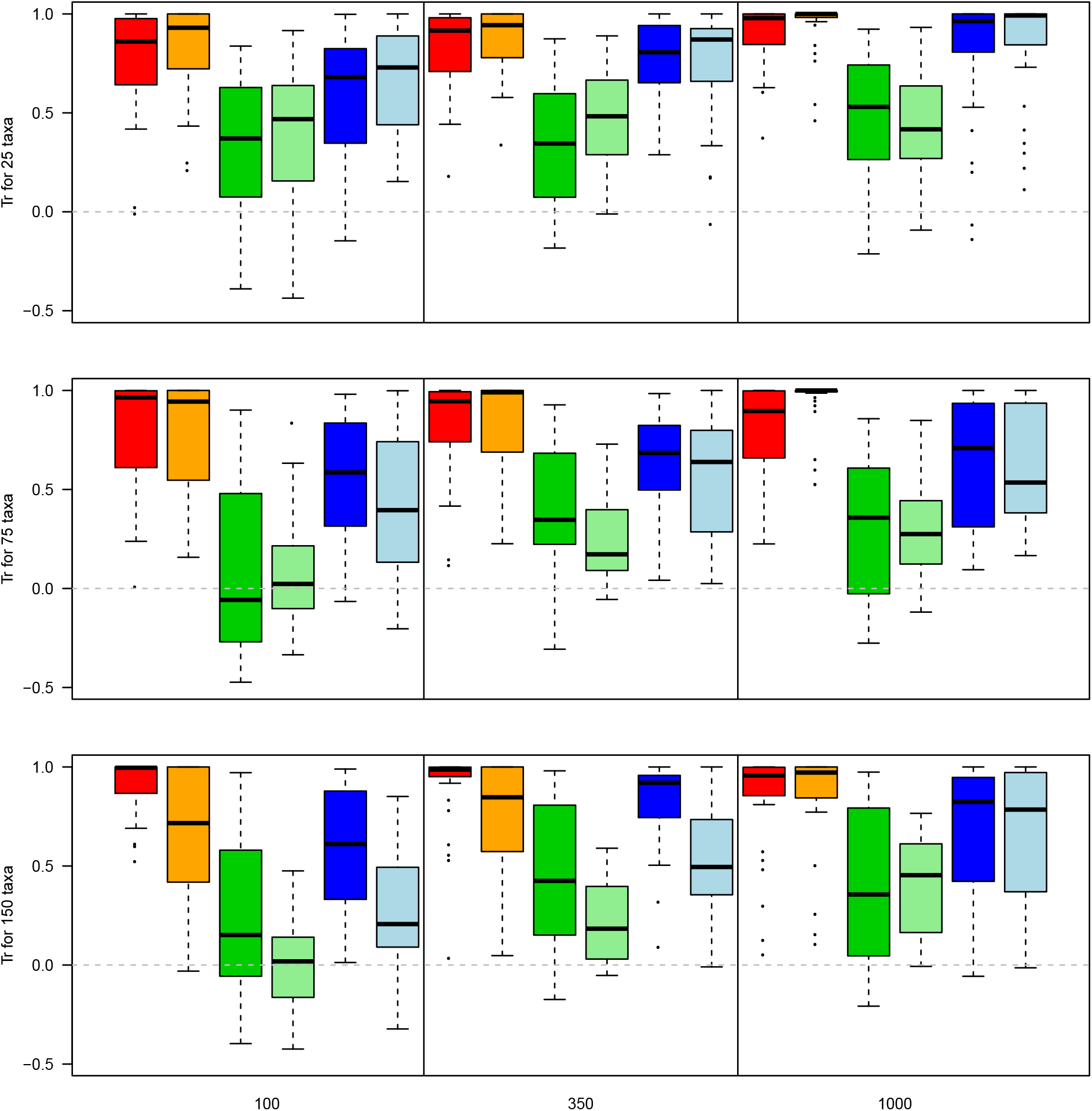
Effect of character difference on recovering the “normal” topology. The axis are identical to figure 3 but y axis represents the Normalised Tree Similarity using Triplets distance.

The metric scores where calculated using the TreeCmp javascript (Bogdanowicz et al., 2012). The measurements where then standardised using the Normalised Tree Similarity metric (*NTS*; i.e. centering the metrics scores using the mean metric score for 1000 pairwise comparisons between random trees with *n* taxa; Bogdanowicz et al., 2012; Guillerme and Cooper, 2016b). When the normalised metric has a score of one it means both trees are identical, when it has a score of zero it means the trees are no more different than expected by chance and when it has a score < 0 the trees are more different than expected by chance. The normalised score for both metrics thus reflects two distinct aspects of tree topology: (1) the Normalised Robinson-Foulds (*NTS*_*RF*_) Similarity reflects the conservation of clades (i.e. a score close to 1 indicates that most clades are identical in both trees); and (2) the Normalised Triplets Similarity (*NTS*_*Tr*_) reflects the position of taxa (i.e. a score close to 1 indicates that most taxa have the same neighbours in both trees).

Because both *NTS*_*RF*_ and *NTS*_*Tr*_ metrics are bounded at one. The residuals of any model based on the *NTS* scores were not normal thus preventing the use of parametric tests for comparisons (see online material https://rawgit.com/TGuillerme/CharactersCorrelation/master/Analysis/02-EffectCorrelationFullResults.html). Similarly, a non-parametric Wilcoxon rank test (Hollander et al., 2013) would be biased in its p-value calculation due to the presence of equal values in the *NTS* distributions (e.g. when multiple trees are equal to the “normal” tree). Therefore, we used a combination of the Wilcoxon rank test with a Bonferonni-Holm corrections (to ensure our significant results were robust to Type I error rate inflation; Holm, 1979) and a simple non-parametric metric for measuring the probability of overlap between two distributions, the Bhattacharyya Coefficient (*BC*; Bhattacharyya, 1943; Guillerme and Cooper, 2016b; Soto et al., 2016). Thus, additionally to the Wilcoxon test results, we considered distribution to be significantly similar if they had an overlap probability > 0.95 and different if they had an overlap probability > 0.05. Comparisons falling between these range can not be designated as strictly similar/different but can still be ranked (e.g. for three distributions A, B, C, if *BC*_(*A*,*B*)_ = 0.15 and *BC*_(*A*,*C*)_ = 0.65, we cannot consider either distribution significantly different or similar but *B* still has a lower probability of being similar to *A* than *C*).

The resulting full simulation was 3.5TB big so is not shared here (though the parameters are). However, the resulting consensus trees on which the topological differences are calculated are available at https://figshare.com/s/7a8fde8eaa39a3d3cf56.

## Results

### Effect of character differences on topology

The overall amount of character difference in a matrix has an effect of the ability to recover the correct topology when maximising character difference leading to the smallest loss in phylogenetic information (median *NTS*_*RF*_ = 0.956 and median *NTS*_*Tr*_ = 0.839) followed by simply randomising the characters (median *NTS*_*RF*_ = 0.762 and median *NTS*_*Tr*_ = 0.628) and minimising the character difference (median *NTS*_*RF*_ = 0.605 and median *NTS*_*Tr*_ = 0.303 - see supplementary material 3 for the full summary statistics). There is a significant difference between all scenarios (maximising, minimising and randomising) with the highest probability of overlap being between maximising and randomising the character difference (Bhattacharrya Coefficient of 0.873 for the *NTS*_*RF*_ and 0.908 for the *NTS*_*Tr*_ - Table 2) and the lowest probability between maximising and minimising the character difference (Bhattacharrya coefficient of 0.573 for the *NTS*_*RF*_ and 0.614 for the *NTS*_*Tr*_ - Table 2)

**Table 2:** Difference between the pooled scenarios. Bhatt.coeff is the Bhattacharrya Coefficient (probability of overlap), the statistic and the p.value are from a non-parametric wilcoxon test (with Bonferonni-Holm correction)

| metric | test | bhatt.coeff | statistic | p.value |
| --- | --- | --- | --- | --- |
| RF | maxi:mini | 0.573 | 356436.000 | 0 |
|  | maxi:rand | 0.873 | 287225.000 | 0 |
|  | mini:rand | 0.856 | 95841.500 | 0 |
| Tr | maxi:mini | 0.614 | 358800.000 | 0 |
|  | maxi:rand | 0.908 | 288223.500 | 0 |
|  | mini:rand | 0.858 | 98507.500 | 0 |

#### Number of characters

This effect of the character difference is not dependent on the number of characters when looking at clade conservation (i.e. *NTS*_*RF*_). The median *NTS*_*RF*_ was similar for 100, 350 and 1000 characters (0.730, 0.745, 0.767 respectively - see supplementary materials 3) with a significant difference only between 100 and 1000 and 350 and 1000 characters (Table 3). The number of characters affects the character difference more in terms of taxon placement for a low number of characters (median *NTS*_*Tr*_ for 100, 350 and 1000 characters equals 0.544, 0.693, 0.799 respectively - see supplementary materials 3) with a significant difference between 100 and 350 or 1000 characters (Table 3). However, these differences have to be contrasted by a very high probability of overlap between each number of characters and metrics (Bhattacharrya Coefficient always > 0.95) suggesting that the significant effects of the number of characters still leads to really similar distributions.

**Table 3:**
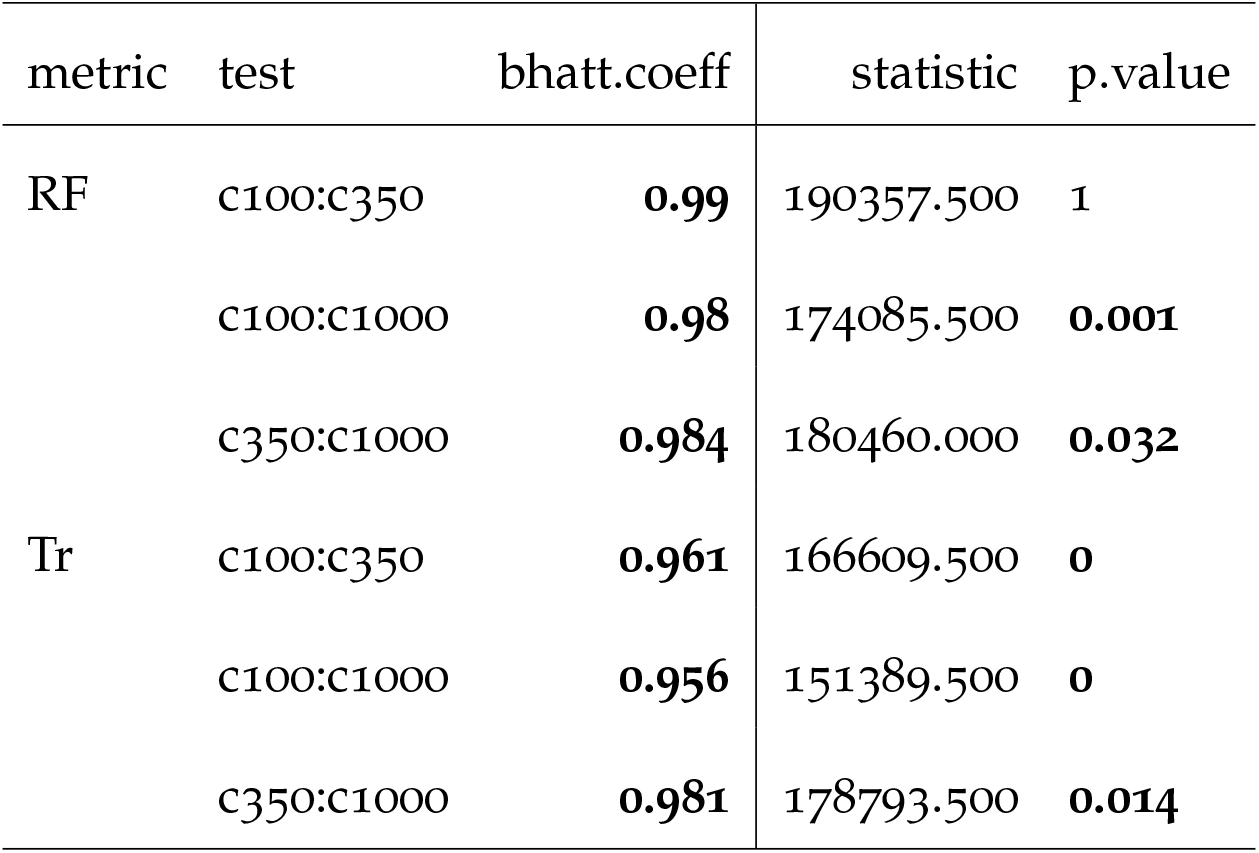
Difference between the pooled number of characters. Bhatt.coeff is the Bhattacharrya Coefficient (probability of overlap), the statistic and the p.value are from a non-parametric wilcoxon test (with Bonferonni-Holm correciton)

| metric | test | bhatt.coeff | statistic | p.value |
| --- | --- | --- | --- | --- |
| RF | c100:c350 | **0.99** | 190357.500 | 1 |
|  | c100:c1000 | **0.98** | 174085.500 | **0.001** |
|  | c350:c1000 | **0.984** | 180460.000 | **0.032** |
| Tr | c100:c350 | **0.961** | 166609.500 | **0** |
|  | c100:c1000 | **0.956** | 151389.500 | **0** |
|  | c350:c1000 | **0.981** | 178793.500 | **0.014** |

#### Number of taxa

Similar to the effect of number of characters on character difference, the number of taxa seems to have only a marginal effect. A low number of taxa (25) resulted in significant differences with both 75 or 150 taxa in both *NTS*_*RF*_ and *NTS*_*Tr*_ but no differences between 75 and 150 taxa (medians for 25, 75 and 150 taxa equals 0.802, 0.76, 0.763 *NTS*_*RF*_ and 0.758, 0.588 and 0.615 *NTS*_*Tr*_ respectively - Table 4 and see supplementary materials 3). Again, however, the significant differences have to be contrasted with still high probabilities of overlaps for each *NTS*_*RF*_ and *NTS*_*Tr*_ distributions for every number of taxa (Table 4).

**Table 4:** Difference between the pooled number of taxa. Bhatt.coeff is the Bhattacharrya Coefficient (probability of overlap), the statistic and the p.value are from a non-parametric wilcoxon test (with Bonferonni-Holm correciton)

| metric | test | bhatt.coeff | statistic | p.value |
| --- | --- | --- | --- | --- |
| RF | t25:t75 | **0.976** | 218421.000 | **0.012** |
|  | t25:t150 | **0.988** | 220529.000 | **0.004** |
|  | t75:t150 | **0.99** | 201037.000 | 1 |
| Tr | t25:t75 | **0.976** | 233282.000 | **0** |
|  | t25:t150 | **0.978** | 227288.000 | **0** |
|  | t75:t150 | **0.992** | 194201.000 | 1 |

### Effect of character differences on the inference method

Regarding the inference method, there is a significant difference in clade conservation between Bayesian and maximum parsimony (Table 5 - median *NTS*_*RF*_ of 0.828 and 0.679 respectively) but not in terms of individual taxon placements (Table 5 - median *NTS*_*Tr*_ of 0.738 and 0.601 respectively).

**Table 5:** Difference between the pooled methods. Bhatt.coeff is the Bhattacharrya Coefficient (probability of overlap), the statistic and the p.value are from a non-parametric wilcoxon test (with Bonferonni-Holm correciton)

| metric | test | bhatt.coef | statistic | p.value |
| --- | --- | --- | --- | --- |
| RF | bayesian:parsimony | 0.891 | 579437.500 | 0 |
| Tr | bayesian:parsimony | **0.984** | 470621.500 | 0.084 |

### Combined effects of taxa, characters and correlation on topology

When looking at the combined effect of each parameter, the “maximised” and “minimised” scenarios are always significantly different with no high probability of overlap for both *NTS*_*RF*_ and *NTS*_*Tr*_ (Wilcoxon rank test p.value < 0.05 and Bhattacharrya Coefficient < 0.95 - see supplementary material 3). The same differences are observed when comparing the “maximised” scenario against the “randomised” one expect for: (1) the Bayesian inference with 25 taxa (with 100, 350 and 1000 characters) and with 75 taxa with 1000 characters for both *NTS*_*RF*_ and *NTS*_*Tr*_; and (2) the maximum parsimony for 25 taxa (with 350 and 1000) characters for both *NTS*_*RF*_ and *NTS*_*Tr*_ and 75 taxa with 100 characters for *NTS*_*Tr*_. Identically, there was always a significant difference between the “minimised” scenario and the “randomised” one was expect for the matrix of 150 taxa and 100 characters under maximum parsimony for *NTS*_*RF*_ and the matrix of 150 and 1000 characters under Bayesian inference for *NTS*_*Tr*_. The full list of comparisons and summary statistics are available in the supplementary materials 3.

## Discussion

### Effect of character differences on topology

As expected, there is a significant effect of the character difference in the ability to recover the correct topology. The character difference metric can be seen as the inverse of character correlation (see Methods): a high character difference approximates a low level of character correlation and vice versa. When characters are correlated, one could expect the matrices to convey a strong (but potentially misleading) phylogenetic signal since every character agrees with each other and conversely, when characters are uncorrelated, one could expect them to convey a weaker phylogenetic signal with a high amount of homoplasy. Intuitively, this would lead the “minimised” character difference scenario to lead to incorrect but consistent trees, the “maximised” scenario to lead to poorly resolved once (really homoplasic trees) and the “randomised” scenario to perform the best at recovering the correct topology. Although the expected results appear to be true for a low character difference scenario, increasing the character difference surprisingly improves the ability to recover the “normal” topology both in terms of clade conservation (*NTS*_*RF*_) and taxa placement (*NTS*_*Tr*_) for both inference methods (especially in bigger matrices; Figs 3 and 4). Furthermore, the trees generated by the “minimised” scenario do not appear better resolved (towards any topology) than the other scenarios (see Supplementary material 3, Figs 3, 4 and 5).

#### Number of characters and taxa

Because of the nature of our simulation protocol, one could expect that the effect of character correlation would have increased with the number of characters (i.e. the more characters available, the more characters are modified in each scenario). Similarly, one could expect the number of taxa to have an effect of the raw ability to recover the “normal” topology (i.e. the more taxa, the more likely taxa are misplaced by chance).

Although we measured a significant difference between “small” and larger matrices (both in terms of number of taxa and characters; Tables 3 and 4), these differences have to be contrasted with the probability of overlap between the results distributions that are always really high between every matrices sizes (Bhattacharrya Coefficients > 0.965 for both the different number of characters and taxa). This suggest that the effect of character correlation on recovering the right topology is independent of the size of the matrix when pooling the data. For the number of characters, this suggests that the overall character difference metric is a good proxy for character correlation as it is independent of the number of characters analysed. Similarly, using the a Normalised Tree Similarity metric (*NTS*) accounts for the fact that topological difference is affected by the sheer number of taxa considered (i.e. we corrected for the expected difference when comparing two random trees with the same number of taxa).

### Effect of character differences on the inference method

When considering the pooled effect of the tree inference method, we only detected a significant difference between the Bayesian and the maximum parsimony trees in terms of clade conservation but none in terms of taxa placement (both using a Wilcoxon test and the Bhattacharrya Coefficient; Table 5). The difference in the ability of each method to recover the “correct” topology has been heavily discussed in the last five years with some indications that Bayesian inference will outperform parsimony when analysing discrete morphological characters alone (Wright and Hillis 2014; O’Reilly et al. 2016; Puttick et al. 2017; although some critics have raised issues with these investigations Spencer and Wilberg 2013; Goloboff et al. 2017). In this study, it is possible that our simulation protocol for generating the characters (favouring slightly more M*k*-based characters rather than HKY ones) could slightly favour Bayesian inference over maximum parsimony, however, our protocol for selecting matrices (i.e. those with in a *CI* < 0.26 in a quick preliminary parsimony search; O’Reilly et al., 2016) could also favour maximum parsimony analysis. It was however not the purpose of this study to compare the overall performance of both methods but rather to measure the effect of character correlation on each of those methods separately.

The differences in performance of the two methods observed here could be due to the inherent mechanisms of each method. For any given topology *T* that was obtained from the “normal” matrix and a matrix with high homoplasy, both methods will generate score differently: (1) in parsimony, the topology will probably be given a bad optimality score (on that implies many changes along the tree) and the optimality criterion (favouring the minimum score) will likely discard the tree. The tree search will thus likely result in a topology island that will not contain the given topology *T*. (2) in Bayesian inference, the topology will also be given a bad optimality score (i.e. low likelihood) although the high homoplasy can be accommodated in the tree through high evolution rates or/and long branches. The rate and the branch length being two parameters among others, the optimality score (the likelihood) will change less drastically than for using parsimony. Furthermore, in Bayesian inference, a reasonable difference in optimality between two topologies (the acceptance) will not necessarily mean that the given topology *T* will be discarded. This difference in both mechanisms could explain why, on average, Bayesian Inference seems better to recover the “normal” topology than maximum parsimony.b

### Distinction between different character correlations

Here we mention three different types of character correlations but evolutionary biologists are mainly interested in the intra-organismal and evolutionary correlations (e.g. in evo-devo Goswami and Janis 2006; or in macroevolution FitzJohn et al. 2014). These two types of correlations can only be studied *a posteriori* with a phylogenetic hypothesis and should not used *a priori* as a criterion to select characters. In other words, intra-organisaml and evolutionary correlation should be studied based on an underlying phylogenetic framework making the correlation induced by data collection (i.e. coding correlation) the only type of correlation that can affect the phylogeny *a priori*. This dichotomy thus creates a trade of between: (1) coding fewer characters (stochastically reducing *a priori* correlation) but making the *a posteriori* correlation more dependent on the coding; and (2) coding more characters (increasing *a priori* dependence) but allowing the *a posteriori* correlation being less dependent on the coding correlations.

It is important to note that the two other sources of character correlation could also be present in our simulations although they were not explicitly modelled: (1) evolutionary correlation is implied by simulating the characters based on Birth-Death trees; and (2) intra-organismal correlation could also be present in the matrices for those characters randomly simulated but sharing similar evolutionary simulation regimes (i.e. creating “modules” of characters). However, the effect of these sources of correlation was out of the scope of this study and would have required *a posteriori* changes to the matrices which are - when using empirical data - at best bad practice and at worth dishonest.

### Limitations

First, simulating evolutionary history is complex. Not only because the models we’re using to infer phylogenies are ever improving (e.g. Heath et al., 2014; Wright et al., 2016) but also because generalising morphological evolution across vastly different organisms is probably impossible (see constrasted discussions from Goloboff et al., 2018; O’Reilly et al., 2018). However, we do not compare the “maximised”, “minimised” and “randomised” to the “true” tree but rather to the “normal” tree. This allows us to reduce the caveats from our simulations on the effect of character correlation since we only compare the simulation end products to each other (the outputs) rather than to the simulation inputs.

Second, measuring and modifying character correlation is difficult. In our simulation protocol we chose to create simulation by duplicating characters in a matrix to maximise or minimise correlation. In biology, this correlation arises from either intra-organismal or evolutionary mechanisms. This could lead to correlations between characters to be more present in some parts of the trees that other (e.g. in the case of inapplicable data Brazeau et al., 2017). However, because of the number of characters, it is actually complex to actually measure their correlation in a biological sense and is still actively discussed in the literature (Russell Lande, 1983; Maddison, 1990; Pagel, 1994; Mark Pagel, 2006; Goswami and Janis, 2006; Goswami and David Polly, 2010; Goswami et al., 2014; Grabowski and Porto, 2016). Additionally, as discussed in the introduction, character correlation can also simply arise by chance due to the discrete coding scheme (i.e. some sets of characters can be highly correlated but effectively describe independent information). Therefore, we made the choice to simplify our simulations by generating character correlation as a stochastic process rather than a biological one.

Third, comparing phylogenetic inference methods is not trivial. As mentioned above, both maximum parsimony and Bayesian inference, although aiming (and often achieving) to infer evolutionary history only have similar outputs and vastly differ in how optimality is measured. But there are also difficulties in summarising both methods with consensus trees OReilly and Donoghue (2017). However, we want to point out again that here we’re no comparing the methods to each other *per se* but rather how they both, individually, react to an increase or decrease of correlated characters.

### Potential applications

Effectively, our simulation protocol bootstraps our data “with bias”. In the “randomised” scenarios the data is simply randomly bootstrapped simply we randomly remove and resample characters (i.e. giving the weight of 0 to some and > 1 to other). However, in the “minimised” and “maximised” scenario, the bootstrapping we remove the characters with the lowest/highest overall character difference. For example, in the “maximised” scenario, we randomly remove some characters that are strongly correlated with other and randomly resample from the left characters.

It is noteworthy to point that in rather small matrices (25 × 100), there was no significant difference in terms of recovering the right topology when maximising or randomising the character differences. Since many discrete morphological matrices are of similar size (Guillerme and Cooper, 2016a) a simple bootstrap re-sampling (i.e. the equivalent of randomising the character differences in our analysis) will be sufficient to obtain a robust topology (*cf.* actively collecting different characters). In matrices with more taxa, however, the “maximised” scenario resulted in better topological recovery than any other scenarios. Applying this kind of bootstraps that maximises character difference by biasing the random sampling could thus results in better resolved trees.

### Conclusion

Correlation between characters can be induced through three main phenomena: intra-organismal relationships, selection-driven covariation or biases in coding the characters yet only the latter can be improved upon to investigate phylogenetic relationships. Useful best practices guidelines (e.g. Brazeau, 2011; Simôes et al., 2017) and algorithms for dealing with different types of character correlations (e.g. for characters hierarchy ?Brazeau et al., 2017) already exist. However, with the regain of popularity in discrete morphological data and the expansion of dataset size (e.g. Ni et al., 2013; O’Leary et al., 2013, with more than 1000 characters each), we can expect the correlation between characters to increase stochastically. Moreover, because phylogenetic inference software are unable to *a priori* differentiate these difference correlations, it is important to understand to what extant topologies can be induced by such bias.

We found that character differences as a proxy for character correlation have a strong effect on recovering the “normal” topology: when character correlation was high (low character differences), the topology was always the furthest away from the “normal” topology. Conversely, when correlation between characters was low, the topology was always the closest to the “normal” topology. These results seem independent on the size of the matrix (number of taxa and/or characters) but can be influenced by the phylogenetic inference method used with Bayesian inference faring better in terms of clade conservation, especially in larger matrices.

However, in modest size matrices (25 taxa; 100 to 350 characters), the effect of actively choosing to minimise character correlation was not more significant than simply bootstrapping the matrix, suggesting that character correlation is more a problem in large discrete morphological matrices. For such matrices, minimising the character correlation (resampling characters < 25% different) or maximising it (> 75%) respectively significantly decreased and increased correct topological recovery compared to randomly resample matrices.

## Data availability, repeatability and reproducibility

The consensus trees are available on figshare at https://figshare.com/s/7a8fde8eaa39a3d3cf56. The simulations are fully replicable following the explanations at https://github.com/TGuillerme/CharactersCorrelation. The post-simulation analysis, tables and figures (reported in this manuscript) are fully reproducible see (https://github.com/TGuillerme/CharactersCorrelation).

## Acknowledgements

Calculations where done using the Imperial College London Cluster Services (doi: 10.14469/hpc/2232). We thank Alberto Pascual Garcia for input in the design of the simulations protocol and Guillermo Herraiz Yebes for helping with the CD metric proof.

